# Analysis of RNA/DNA Hybrid Profiles in Blood From Autistic Patients Reveals Differences in mRNA and non-coding RNA Regions

**DOI:** 10.1101/2024.10.18.618986

**Authors:** Leila Kianmehr, Kasra MokhtarZadeh, Zeynep Yilmaz, Ecmel Mehmetbeyoglu Duman, Serpil Taheri, Minoo Rassoulzadegan

## Abstract

DNA/RNA hybrids are present in normal cellular physiology, but when misregulated in the genome, they can contribute to human pathologies such as neurodevelopmental disorders and cancer. These hybrid structures are often abundant in repetitive genomic sequences such as telomeres and centromeres, contributing to evolutionary conserved phenomena. However, excessive accumulation can lead to DNA damage and genomic instability.

Studies of autism spectrum disorders (ASDs) have revealed genome-wide disruptions, including mutations and significant deregulation of gene expression. Surprisingly, the common underlying factors remain unidentified. ASDs, which are of hereditary origin and more prevalent among boys, manifest themselves during development through disorders of various origins.

In this study, we used whole-genome transcriptomic analysis of DNA/RNA hybrids from blood samples of autistic patients and healthy controls. we identified differences between numerous hybrid genomic regions with varying sequence compositions. Additionally, we observed local specificity in genomic regions, where RNA could either inhibit or enhance transcription. Our analysis of short RNAs hybridized with DNA revealed a striking example of transcriptional activity, where each gene is stably transcribed. We propose that non-coding RNAs originating from exons, introns or intergenic regions bind to DNA as hybrids (DNA/RNA) and influence gene expression. The specific levels of these short RNAs may regulate their own transcription, likely affecting cell fate.

## Introduction

During RNA trancription, one strand of DNA is displaced, and the newly synthesized RNA forms a hybrid DNA/RNA structure. This occurs continuously; some RNA strands remain in nucleus, some are transferred to the cytoplasm after maturation, and some maintain as hybrids with DNA until they are digested by RNaseH before replication. This process results in displaced DNA helices that form DNA/RNA hybrids. These hybrids regions are present in normal cellular physiology across several genomic areas, but changes in their amount and position are associated with diseases (Crossley et al., 2019; García-Muse & Aguilera, 2019; Marnef & Legube, 2021; Niehrs & Luke, 2020; Petermann et al., 2022). DNA/RNA Hybrids are abundant in specific repetitive sequences such as telomeres (Arora et al., 2014; Feretzaki et al., 2020), and centromeres (Kabeche et al., 2018; Y. Liu et al., 2021), contributing to evolutionarily conserved phenomena (Chen et al., 2017; Ginno et al., 2012, 2013; Hartono et al., 2018; Sanz et al., 2016; Skourti-Stathaki et al., 2011; Xu et al., 2017). Excessive hybrid formation in repeated motifs within postmitotic neurons has the potential to enhance transcription, leading to DNA damage and genomic instability, and potentially contributing to neurodevelopment and neurodegenerative disorders. Mutation in DNA repair genes, such as ATM, SETX, are associated with DNA damage in neurons. Notably, SETX, which encodes the Senataxin protein involved in resolving hybrids, has been linked to this process (Skourti-Stathaki et al., 2011).

Several neurodegenerative disorders, including ALS, dementia, Fredrich Ataxia, and Fragile-X are associated with expansion repeats prone to forming R-loops (Groh et al., 2014; Haeusler et al., 2014; Richard & Manley, 2017). In these conditions, trinucleotide repeats coincide with variable transcription rates. Hybrids at expanded repeats are correlated with impaired Pol II function and gene silencing, suggesting that hybrids contribute to pathology by posing blocks to gene expression (Groh et al., 2014; Haeusler et al., 2014; Walker et al., 2017). In other disorders, such asnnAicardi–Goutières syndrome, mutations in genes involved in hybrids resolution (e.g., *TREX1* or in *RNASEH2*) lead to global increases in DNA/RNA hybrids (Lima et al., 2016; Mackenzie et al., 2016; Coquel et al., 2018).

It is well-stablished that RNA plays a regulatory role in gene expression at transcriptional, post-transcriptional, and translational levels in eukaryotes (Mattick et al., 2023). Hybrid formation modulates transcription and recombination, suggesting that these structures likely have regulatory functions through their abundance and transient positions in the genome. For example, TERRA (Telomeric Repeat-containing RNA) forms stable DNA/RNA structure at telomeres, regulating telomere length surveillance. Short telomeres accumulate TERRA (Feretzaki et al., 2020). We have developed a new method to measure TERRA amounts engaged in hybrids at chromosome ends (Rassoulzadegan et al., 2020).

Despite the increasing prevalence of autism spectrum disorder (ASD) and the significant research conducted (Li et al., 2023; Yasuda et al., 2023) surprisingly little is known about its mechanism and aetiology. ASD diagnosis is based on abnormal behavior, including impairments in social interactions, communication, and repetitive stereotypic behaviors (Iakoucheva et al., 2019). Although genetics and environmental factors are implicated in altered brain development and neural activity (Auerbach et al., 2011; L. Liu et al., 2014), no consistent genetic association thas been identified, as evidenced by recent large-scale exome sequencing analysis (Satterstrom et al., 2020). This lack of a uniform genetic signature highlights the inherent heterogeneity within ASD, which is also reflected in transcription levels (Masi et al., 2017).

Recently, we reported progressive heritable downregulation of a group of miRNAs in autistic patients and their families (Ozkul et al., 2020). Gene silencing in various organisms is often linked to the expression of specific RNA in cis or trans (Tufarelli, 2006).

We are interested in detecting whether hybrid regions in the genome of patients with autism differ from those in controls. Current methods for detecting DNA/RNA hybrids have limitations and often produce inconsistent results (Bou-Nader et al., 2022; Gibbons & Aune, 2020; Halász et al., 2017; Sanz & Chédin, 2019). To address this, we recently developed an antibody-independent method to identify genomic regions with hybrid structures, which allows profiling of nascent transcripts (Kianmehr et al., 2019; Mehmetbeyoglu et al., 2022; Rassoulzadegan et al., 2020). In this study, we applied the same strategy to identify DNA/RNA hybrids from blood samples of patients with autism and healthy controls.

Combining bioinformatics with molecular biology techniques, we conducted transcriptome profiling of autistic patients and healthy controls. We focused on transcriptome-guided assembly on DNA/RNA hybrids using RNA-seq data from autistic patients. We then performed differential gene expression analysis to characterize both known and novel transcripts bound to DNA in DNA/RNA hybrids.

In summary, our transcriptome-wide analysis reveals alterations in the profile of DNA/RNA hybrids in mRNA and ncRNA regions of autistic patients. These changes in hybrids provide insights into how the transcriptome is altered in ASD.

## Material and Methods

### DNA/RNA hybrid preparation

DNA/RNA hybrids have traditionally been detected using DRIP-assay or enzymatic recognition by RNaseH. However, the specificity of the S9.6 antibody for accurate R-loop quantification and mapping has recently been questioned (Bou-Nader et al., 2022). Additionally, the efficiency of chromatin fragmentation by restriction enzymes in DRIP-related approaches may affect the resolution of R-loop mapping (Halász et al., 2017). To address these issues, we developed a direct strategy for investigating DNA/RNA hybrids without relying on antibodies (Kianmehr et al., 2019; Rassoulzadegan, Sharifi-Zarchi, & Kianmehr, 2021, Mehmetbeyoglu et al., 2022). This method modifies classical TRIzol-chloroform extraction protocol, allowing for the isolation of DNA/RNA hybrids from total RNA without antigenic or RNaseH recognition, thereby avoiding potential artifacts.

After the enzymatic removal of proteins, total nucleic acids were isolated, and DNA and RNA were fractionated using the standard TRIzol^TM^ protocol (Rio et al., 2010). The nucleic acid fractions from the aqueous phase (RNA) and chloroform-water interphase were ethanol-precipitated separately. These fractions were then re-fractionated through Zymo-Spin^TM^ columns (ZYMO-RESEARCH CORP, Irvine, CA, USA) and eluted or further purified via chloroform extraction.

Ethanol precipitated interface materials underwent overnight incubation at 56°C in a buffer containing 20 mM Tris (pH 8), 50 mM EDTA, 0.5% SDS, 20 µM dithiothreitol, and 400 µg/mL Proteinase K. Free RNA was separated, and DNA/RNA hybrids were further purified following DNase digestion and additional purification using Zymo-Spin^TM^ columns (ZYMO-RESEARCH CORP, Irvine, CA, USA). Extracts were treated with RNase (to remove all RNAs), RNaseH (to remove only DNA-RNA hybrids), and DNase (to remove all DNA fragments). Following the enzymatic removal of proteins, total nucleic acids were fractionated into RNA and DNA-bound RNA.

### RNA Sequencing (RNA-seq)

Blood samples were collected from six autistic patients and healthy controls at the Erciyes University School of Medicine Hospital, Kayseri, Turkey. This study received approval from the Ethics Committee of the Erciyes University School of Medicine (Committee No: 2011/10, approval date: 09-20-2011). The diagnosis was made according to the criteria of the Diagnostic and Statistical Manual, Fourth and Fifth Edition, Text Revision (DSM-IV-TR; American Psychiatric Association, 2000 and DSM-V; American Psychiatric Association, 2013) using the Childhood Autism Rating Scale (CARS) (Schopler et al., 1980). All subjects were carefully screened for signs of infection, and subjects with acute illness were excluded.

RNA was recovered from blood samples after DNase digestion with amounts from 10 to 100 ng. High-throughput sequencing was performed on the Illumina HiSeq 2500 or MiSeq platform (Eurofins Medigenomix GmbH, Ebersberg, Germany) (Rassoulzadegan et al., 2021). RNA libraries were prepared from biological replicates of DNA-bound RNA fractions from the autistic patients.

### Bioinformatics analysis

Quality control of the raw sequencing data was conducted using FastQC version 0.11.9. low-quality reads and adapter sequences were trimmed using Trimmomatic version 0.39 (Bolger et al., 2014) which involved removing 12 nucleotides from the 5′ end of each read and bases with a Phred score lower than 20 from both 5′ and 3′ ends. A minimum threshold of 35 bps was set for the reads after trimming. Trimmed reads were aligned to the reference genome (hg38) using Hisat2 (M. Pertea et al., 2016) with the command: ‘hisat2 –dta –rna-strandness’. Samtools was employed for sorting and indexing the alignment files.

### Transcriptome Assembly

To eliminate potential ribosomal RNA, RSeQC (Wang et al., 2012) version 4.0.0 was utilized. Genome-guided transcriptome assembly was performed using StringTie (Kovaka et al., 2019) version 2.1.1 using an annotation file in GFF3 format (*gencode.v39.chr_patch_hapl_scaff.annotation.gff3)* obtained from (https://www.gencodegenes.org/human). Individual sample transcriptome were created, and merged using StringTie merge command to generate a reference transcriptome assembly for quantification. Strand-specific settings were applied in both alignment and assembly steps. Gene expression quantification was conducted using HTSeq-count (Anders et al., 2015) version 0.11.2. For visualization, Integrative genome viewer (IGV) (Thorvaldsdóttir et al., 2013) version 2.12.0 was utilized. Reads were cross-sample normalized to 10^7^ reads per sample for comparative visualization.

### Differential Gene expression analysis

Differential gene expression analysis was performed using DESeq2 (Love et al., 2014) version 1.36.0. P-values were adjusted for multiple comparisons using the Benjamini and Hutchberg method. Genes with an absolute log2 fold change > 1 and an adjusted p-value < 0.05 were considered differentially expressed. Data visualization was conducted using R packages, including ggplot2 (Wilkinson, 2011) version 3.3.6 and pheatmap (Kolde, 2022) version 1.0.12.. Gene ontology (GO) and KEGG pathway over-representation analyses were performed by Cluster Profiler (Wu et al., 2021) version 4.4.4.

### Detection of novel lncRNAs

Novel transcripts were identified using GffCompare (G. Pertea & Pertea, 2020) version 0.12.6 focusing on intergenic and antisense transcripts with a minimum length of 200 nucleotides, which were selected for coding potential assessment. SeqKit (Shen et al., 2016) and GffRead (G. Pertea & Pertea, 2020) were used to extract the nucleotide sequences. The coding potential of each transcript was assessed with CPAT (Wang et al., 2013), selecting the highest potential across all open reading frames (ORFs). Transcripts with coding potential below 0.364 were classified as novel lncRNAs.

### Peak finding

HOMER (Heinz et al., 2010) version 4.11 was employed to identify significant enriched peaks (DNA/RNA hybrid accumulation sites), utilizing the ‘findpeaks *–o auto’* and ‘*annotatePeaks.pl’* commands with a false discovery rate (FDR) threshold of 0.001. Annotated positions included promoter-TSS (Transcription Start Site), 5’ UTR (Untranslated Region), exon, intron, 3’ UTR (Untranslated Region), TTS (Transcription Termination Site), intergenic, and non-coding regions, based on hg38.

### qRT-PCR

Total RNA samples were reverse transcribed into complementary DNA (cDNA) using the Evoscript Universal cDNA Master Kit (Roche, Mannheim, Germany, Cat No: 07912439001) in final reaction volumes of 20 µL, adhering to the manufacturer’s protocol. The cDNA samples were stored at −80°C until subjected to quantitative PCR (Q-PCR) analysis.

Quantitative real-time PCR (RT-PCR) validation was performed on six selected significant differential transcripts using the LightCycler 480 II high-throughput Real-Time PCR system (Roche, Mannheim, Germany). The cDNA samples were diluted in a 1:5 ratio with nuclease-free water. SYBR Green Master (Roche, Mannheim, Germany, Cat No: 04707516001) was employed to quantify transcript levels of long non-coding RNA (lncRNA) SLC12A5-AS1 and RN7SK, as well as *SLC16A3, NLGN3, SMARCC2, ADAMTSL4,* and the reference gene human beta-actin (ACTB).

The reaction mix was prepared according to the manufacturer’s guidelines, with *ACTB*. Changes in gene expression were calculated using the 2^−ΔΔCt^ relative quantification method across all groups (Livak & Schmittgen, 2001).

## Results

Autism, as an inherited disorder, cannot be explained solely by genetics. Faced with the challenge of explaining the origin of the variation in autism, we aimed to establish a robust transcriptome analysis. Specifically, we sought transcriptional signals, such as changes in transcription rates, that could be influenced by environmental factors. Recently, we developed a method that allows reproducible analysis of emerging transcripts. RNA extracted in the form of hybrids with DNA produces highly consistent results, likely due to protection of RNA during the extraction process when hybridized with DNA. In this study, we identify a fraction of RNA signals that may be modified by environmental factors and transmitted through non-Mendelian inheritance. Fraction of DNA/RNA hybrids were extracted from blood samples of six autism patients and six healthy control, yielding a total of approximately ∼ 350 million sequence reads across 12 biological replicates and four technical replicates for each patient and control. The extraction of DNA/RNA hybrid fractions followed (see Methods) we the same method previously described (Kianmehr et al., 2019; Rassoulzadegan et al., 2020). Then, RNA-sequencing libraries were generated from these fractions and 3.5 × 10^8^ total paired-end reads were generated. After quality control, five samples (three patients and two controls) were selected for final analysis.

Genomic mapping of RNA-seq reads was performed using the human GRCh38 genome assembly, summarized ine the pipeline schematic in Figure 1a. Only uniquely mapped paired-ends reads were used for genome-guided transcriptomes assembly. Potential ribosomal RNAs and reads not aligned to the reference genome were filtered out using the RseQC tool. The remaining reads were submitted to the StringTie transcriptome assembly program (Kovaka et al., 2019).

**Figure 1:**
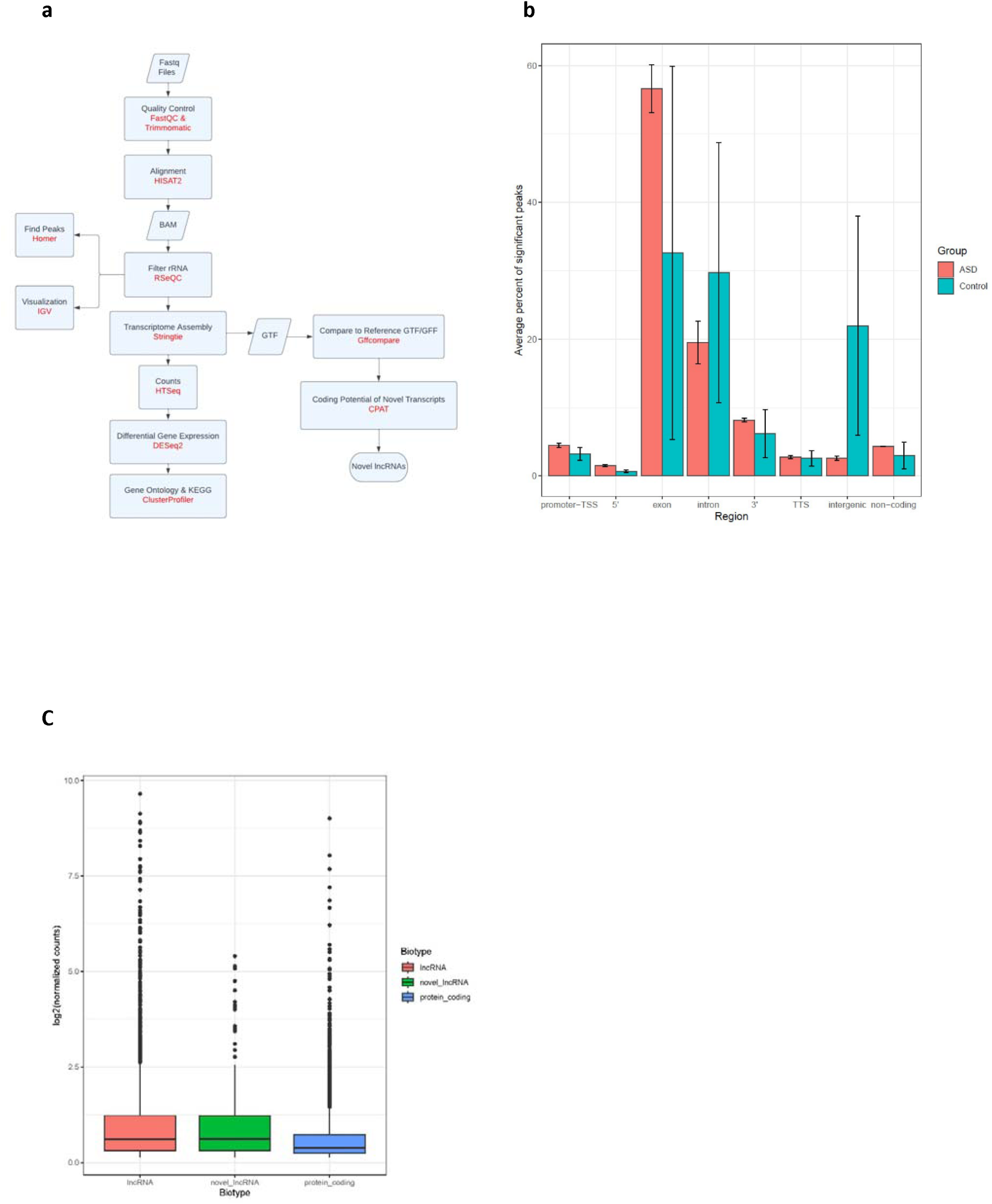
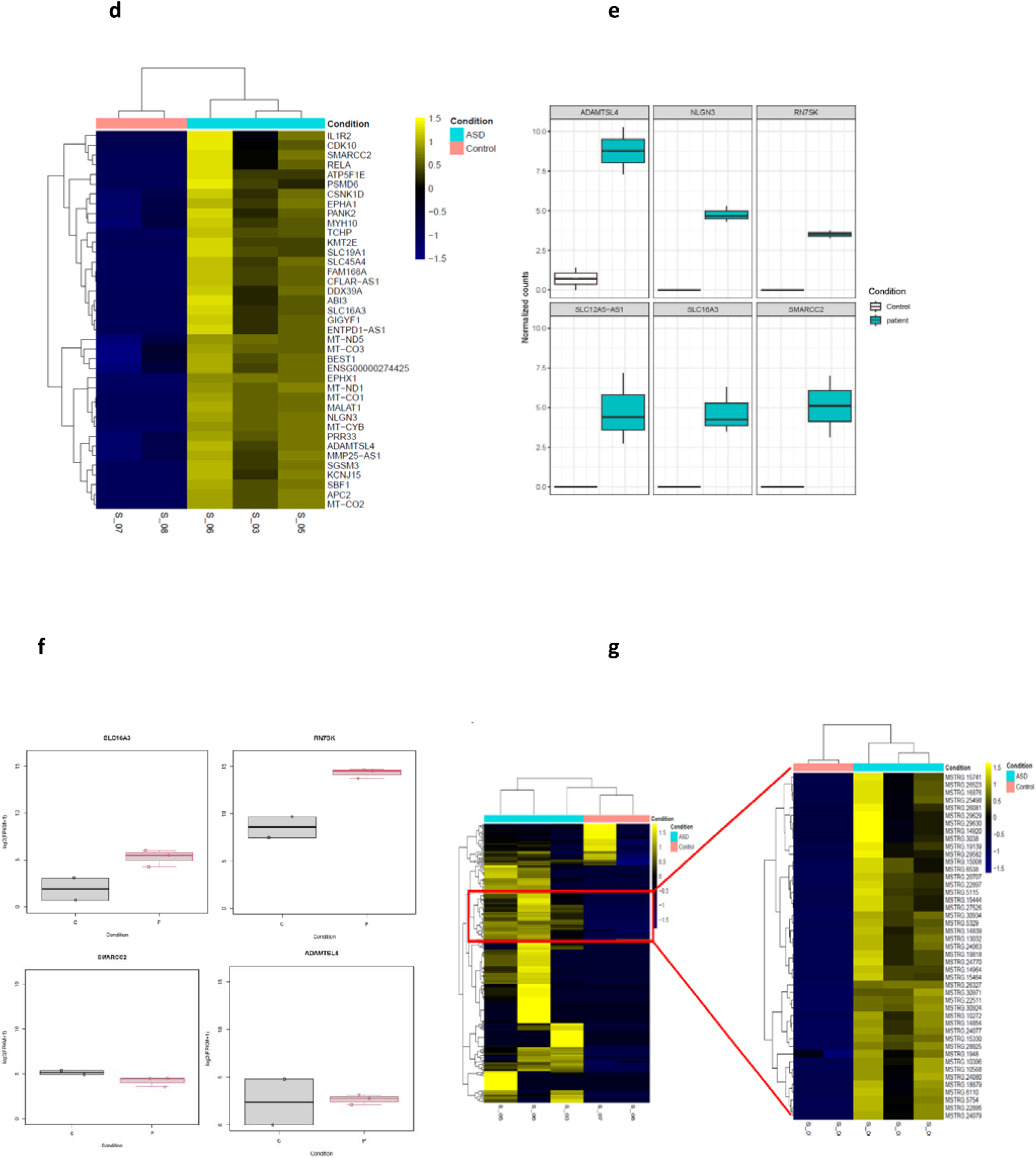
**a,** Schematic representation of the RNA-seq data analysis pipeline. **b,** The average percentage of significant peaks across different genomic regions of DNA/RNA hybrids in autistic patients and healthy controls. The annotated positions include key regions such as promoter-TSS (Transcription Start Site), 5’ UTR (Untranslated Region), exon, intron, 3’ UTR (Untranslated Region), TTS (Transcription Termination Site), intergenic, and non-coding regions. Notably, the exonic regions contain the majority of significant peaks. A false discovery rate (FDR) threshold of 0.001 was applied to identify significant peaks. Error bars represent mean ± SEM. **c,** Expression levels of lncRNAs, novel lncRNAs and protein-coding transcripts across all samples. **d,** Heatmap showing the most significant differentially expressed transcripts comparing autistic patients (ASD) to healthy control (Control). Yellow bars indicate high expression level (LFC>2, *P* value < 0.05), while and blue bars indicate low expression levels (LFC < −2, *P* value < 0.05). Raw Z-scores of log-normalized counts were used to generate the heatmap. e, Normalized counts of significant differentially expressed transcripts (ADAMTSL4, NLGN3, RN7SK, SLC12A5-AS1, SLC16A3, and SMARCC2) in autistic patients compared to healthy controls. f, Log2(FPKM+1) of significant differentially expressed transcripts (ADAMTSL4, NLGN3, RN7SK, SLC12A5-AS1, SLC16A3, and SMARCC2) in autistic patients versus healthy controls **g,** Heatmap showing the expression of novel lncRNAs across all samples (left). Hierarchical clustering revealed a gene module containing 45 novel lncRNAs (red box) which showed an accumulation of DNA/RNA hybrids in ASD samples (right).

### 1 Genome-Guided Transcriptome Assembly Of DNA/RNA Hybrids Revealed Unannotated Transcripts

We identified 278,300 transcripts from the assembly, distributed across 68,487 DNA/RNA hybrid loci. This assembly was performed using the ‘**stringtie –rf**’ command.

Transcripts generated by *StringTie 2.1.1* consistently returned high support scores, indicating the high-quality (high sensitivity and accuracy) of the transcripts returned from the transcriptome assembly (Supplementary File, Table 1).

Our objective was to quantify the transcriptome characteristics and changes of DNA/RNA hybrids in autism blood cells in different genomic regions.

A reference gene annotation file (GTF) containing 263,306 transcripts at 63,463 loci across the genome was provided to *StringTie*. Including this reference file significantly improved the recovery of transcript structure. As a result, we identified 9,501 new exons and 2930 new introns with high sensitivity and accuracy in 5,196 loci (Supplementary File, Table1).

Of the 278,300 transcripts identified, 49% percent were protein-coding, 36% were long noncoding RNAs, and 15% were pseudogenes transcripts (supplementary Figure 1).

### 2 Peak Finding Reveals Distinct Regional Transcripts in Autism Blood Cells

We performed a genome-wide distribution analysis to assess whether specific genomic regions of DNA/RNA hybrids in blood cells of autism patients were associated with altered transcription. Using Homer (Heinz et al., 2010), we analyzed genomic regions based on the GRCh38 genome assembly, including promoter-TSS, 5’ UTR, exon, intron, 3’ UTR, TTS, intergenic, and non-coding regions.

The genome-wide distribution revealed a significant accumulation of DNA/RNA hybrids in exonic and intronic regions (Figure 1. b). In autism samples, more than half of the hybrids were located in exonic regions. Control samples exhibited a similar pattern but had a higher proportion of hybrids in intergenic regions.

### 3 Differential lncRNA Expression Profiles in Autism Patients

Both novel and known noncoding genes showed slightly higher expression levels than annotated protein-coding genes (Figure. 1 c). After filtering low-expressed transcripts, we detected 13,305 known transcripts across all samples. (see Supplementary file-Figure 2a, b).

Differential expression analysis using DESeq2 revealed 702 transcripts that were differentially expressed, with 301 transcripts significantly upregulated, showing a higher density of DNA/RNA hybrids in ASD patients (LFC> 2, adjusted *P* value < 0.05), while 401 transcripts were significantly downregulated (LFC<-2, adjusted *p* value <0.05) (refer to Supplementary-figure 5). The most significant up-regulated transcripts are illustrated based on log2 fold change and baseline mean (Figure 1d, Supplementary Figure 2 a,b). Some of these transcript, including protein coding transcripts (*ADAMTSL4, NLGN3, SLC16A3, SMARCC2*), and lncRNAs (*RN7SK, SLC12A5-AS1*) are demonstrated (Fig 1e). Among them, *SLC16A3, RN7SK,* and *ADAMTSL4* have higher expression (log2FPKM+1) in patients’ samples than healthy controls (Figure 1f). The expression levels of known transcripts (lncRNA, mRNA, protein-coding, snRNA, snoRNA) were categorized by different biotypes (Supplementary file-Figure 3b).

To investigate novel lncRNAs from selected unannotated transcripts associated with DNA/RNA hybrids (antisense and intergenic regions) in autism patients, we considered only transcripts with a minimum length of 200 nucleotides. In total, 1755 transcripts were selected. The Coding Potential Assessment Tool (CPAT), a logistic regression model that predicts coding probability from nucleotide sequences (Wang et al., 2013) (Supplementary Figure 3a,b), was then applied, which predicted that the majority of these transcripts were noncoding. The final protein-coding probability for each transcript was determined by selecting the highest probability among all open reading frames, using a threshold of 0.364. Analysis of exon distribution in novel lncRNAs revelaed that 58.1% of these lncRNAs consisted of a single exon, which less than 40% contained two or more exons (36.1%) (Supplementary Figure 4). From the 1755 transcripts analyzed, 707 transcripts were classified as protein-coding (coding probability > 0.364). While, 1,048 transcripts were identified as novel lncRNAs (coding probability < 0.364) (Supplementary Figure 3a).

None of these noncoding genes were present in the Gencode v39 reference transcriptome annotation, thus classifying them as novel transcripts. Notably, none of the novel lncRNAs found in the differentially expression results of this study. However, principal component analysis (PCA) of the novel lncRNAs revealed variation between autism patients and healthy controls (see Supplementary Figure 5). Highly enriched lncRNAs, which demonstrated a contiguous accumulation of DNA/RNA hybrids in ASD samples compared to healthy controls are shown in Figure 1h. Hierarchical clustering of these novel lncRNAs revealed a module containing 45 novel lncRNAs that were consistently accumulated across all patient samples (Figure 1g).

#### Integrative Genomics Viewer (**IGV) Visualization :**

We compared the differentially expressed transcripts with the SFARI database (https://gene.sfari.org/), a comprehensive resource on the genetic causes of autism. From our gene list, we selected transcripts reported in the SFARI database that are related to autism. To further compare the mRNA expression levels of these selected transcripts across different tissues, we used the GTEx portal (https://gtexportal.org/home/) (Supplementary Figure 6). As a result, we identified 10 transcripts strongly associated with ASD, including *GIGYF1*, *SMARCC2*, *EPHA1*, *IL1R2*, *KCNJ15*, *KMT2E*, *MYH10*, *NLGN3*, *SBF1*, and *SGSM3* (Supplementary Table2). Visualization of key genes, such as *SLC16A3* and *GIGYF1,* using IGV revealed significant signals in ASD samples compared to healthy controls (Figure 2). Additionally, we observed that other transcripts, strongly associated with DNA/RNA hybrids, such as RN7SK and mitochondrial genes (MT-ND5 and MT-CYB) —also exhibited significant signals in ASD samples compared to healthy controls (Figure 2).

**Figure 2.**
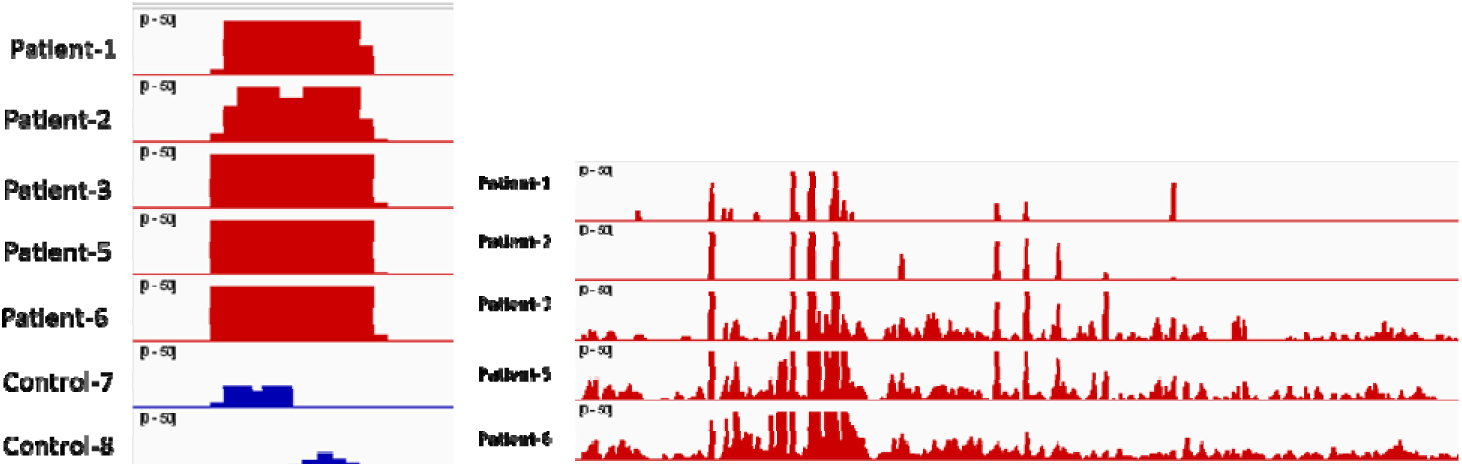
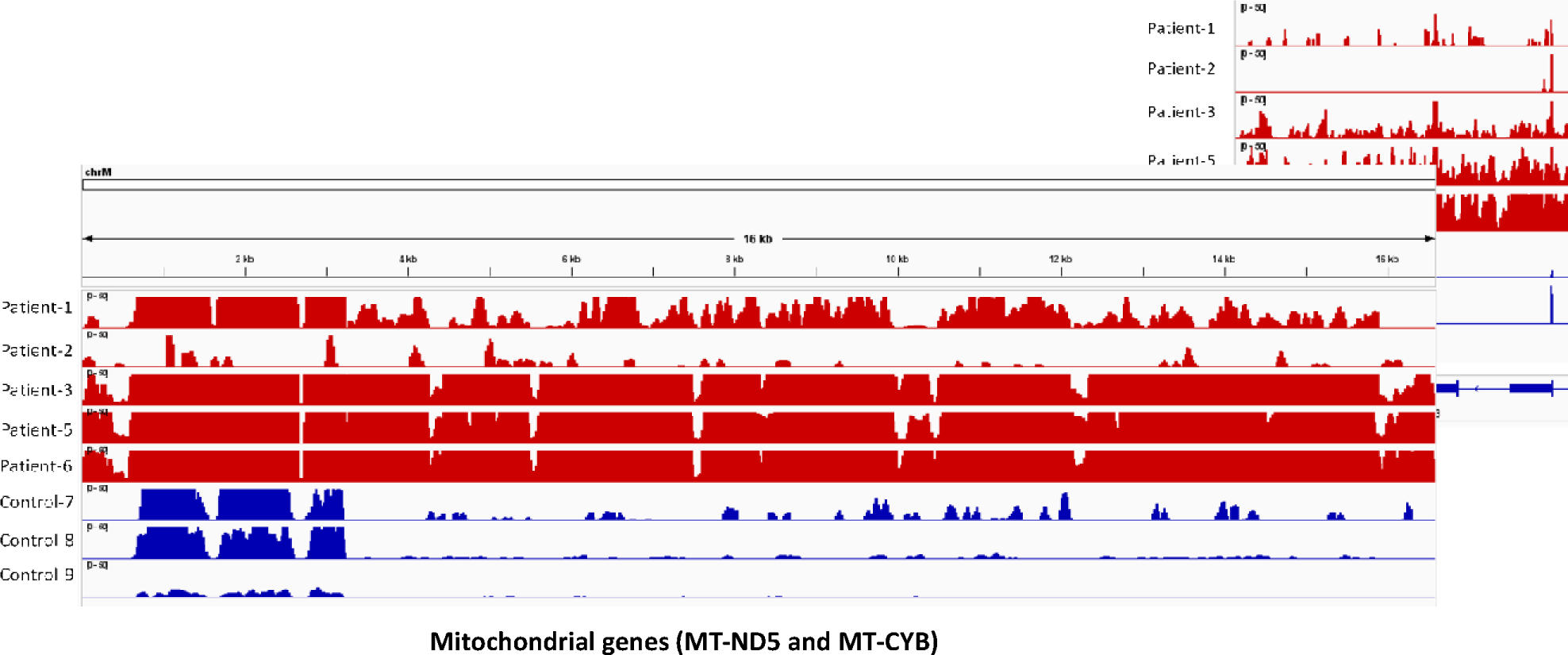
Visualization of ASD risk genes, including *RN7SK* (chr6:52,995,773-52,996,031), *GIGYF1* and *SLC16A3* and mitochondrial genes (*MT-ND5* and *MT-CYB*), shows prominent signals in ASD (red tracks) compared to healthy controls (blue tracks). The green arrow displays the loci selected for qRT-PCR validation.

#### Gene Ontology and KEGG Pathways Analysis

Gene ontology (GO) analysis and KEGG pathway analysis were performed on transcripts differentially detected in DNA/RNA hybrids from autistic patients (log2-fold change > 1, *p* value < 0.05). These analyses were conducted with the ClusterProfiler 4.4.4 package and org.Hs.eg.db (Genome wide annotation for Human) version 3.15.0.

The most significant GO terms enriched for Biological Process (BP), Molecular function (MF), and Cellular Component (CC) were identified (Supplementary Figure 7). The results showed the majority of RNAs in autistic patients were primarily associated with mitochondrial electron transport chain complexes and cellular respiration. Notably enriched terms included aerobic respiration, electron transport coupled to mitochondrial ATP synthesis, mitochondrial respirasome, inner mitochondrial membrane protein complex, and transmembrane transporter activity. These findings emphasize the crucial role of <u>cellular respiration</u> in ASD.

In addition, KEGG pathway over-representation analysis revealed pathways linked to neurodegeneration and neurodegenerative diseases, such as Alzheimer’s disease, Parkinson’s disease, amyotrophic lateral sclerosis, and Huntington’s disease (Supplementary Figure 8).

Further investigation of genes involved in enriched GO terms (Supplementary Figure 9) highlighted key genes like *MT-ND1* and *MT-ND5* of electron transport chain (ETC) complex I, *CYTB* (*MT-CYB*) from ETC complex III, *COX1* (*MT-CO1*), *COX2* (*MT-CO2*), and *COX3* (*MT-CO3*) from ETC complex IV, and *ATP5F1E* from ETC complex V.

#### Validation of RNA-seq results by Quantitative Real-Time PCR Analysis (qRT-PCR)

RNA-seq analysis, which examines the entire transcriptome of DNA-associated RNAs through deep sequencing identified six transcripts significantly differentially expressed in autism patients. These findings were validated through qRT-PCR, which confirmed the significant differential expression of these transcripts in autism patients compared to healthy controls.

DNA/RNA hybrids (D fraction) and total RNA (free RNA fraction) were isolated from blood samples of patients with autism and healthy controls. The results show that the selected transcripts had higher expression level (LFC>2) in the D fractions, suggesting their involvement in autism, as supported by the SFARI database. These transcripts were further quantified using qRT-PCR to validate the RNA-seq findings.

The relative expression level of six transcripts (*NLGN3, SMARCC2, ADAMTSL4, SLC16A3, RN7SK, SLC12A5-AS1*) in total RNA and DNA-associated RNA fractions from autism patients and healthy controls were measured by qRT-PCR. Transcript levels were normalized to *GAPDH* as a reference gene with the primers listed in Supplemental Table 3.

The results revealed that most transcripts were more highly expressed in D fractions compared to the free RNA (R) fractions, consistent with the RNA-seq analysis. Among these, *RN7SK* and *SMARCC2* showed statistically significant difference (p < 0.05). In the free-RNA fractions (total RNAs), a significant difference (means± SEM) were also observed for *ADAMTSL4* and *SLC16A3* (p < 0.05). While NLGN3, SMARCC2 and RN7SK showed higher expression in the total RNA of autism patients compared to healthy controls, the difference were not statistically significant (Figures 3 & 4).

**Figure 3.**
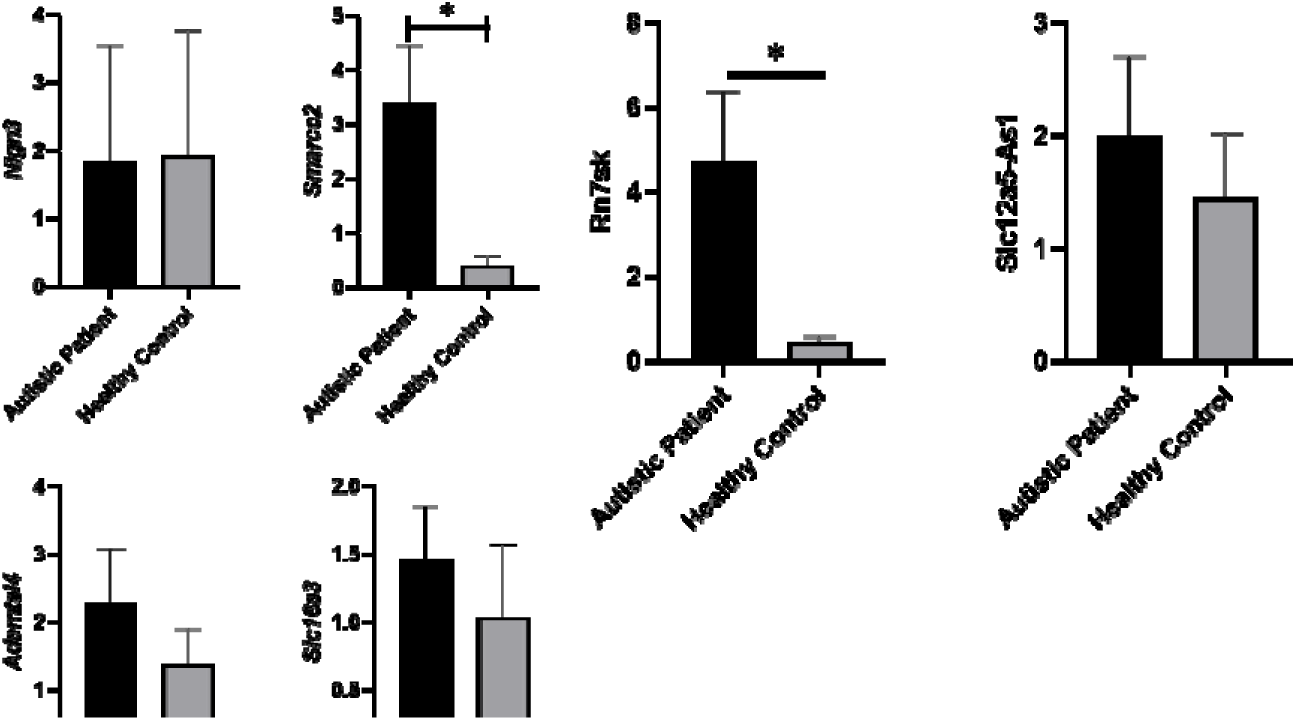
Relative expression levels of six candidate transcripts, Nlgn3, Smarcc2, Adamtsl4, Slc16a3 (protein-coding), and Rn7sk, Slc2a5-As1 (lncRNAs) in DNA/RNA hybrids (D) vs. GAPDH (log2). The levels in each sample were normalized to that of GAPDH (log2) (internal control). Data are presented as means± SEM. Group mean comparisons are performed using Student’s t-test.

**Figure 4.**
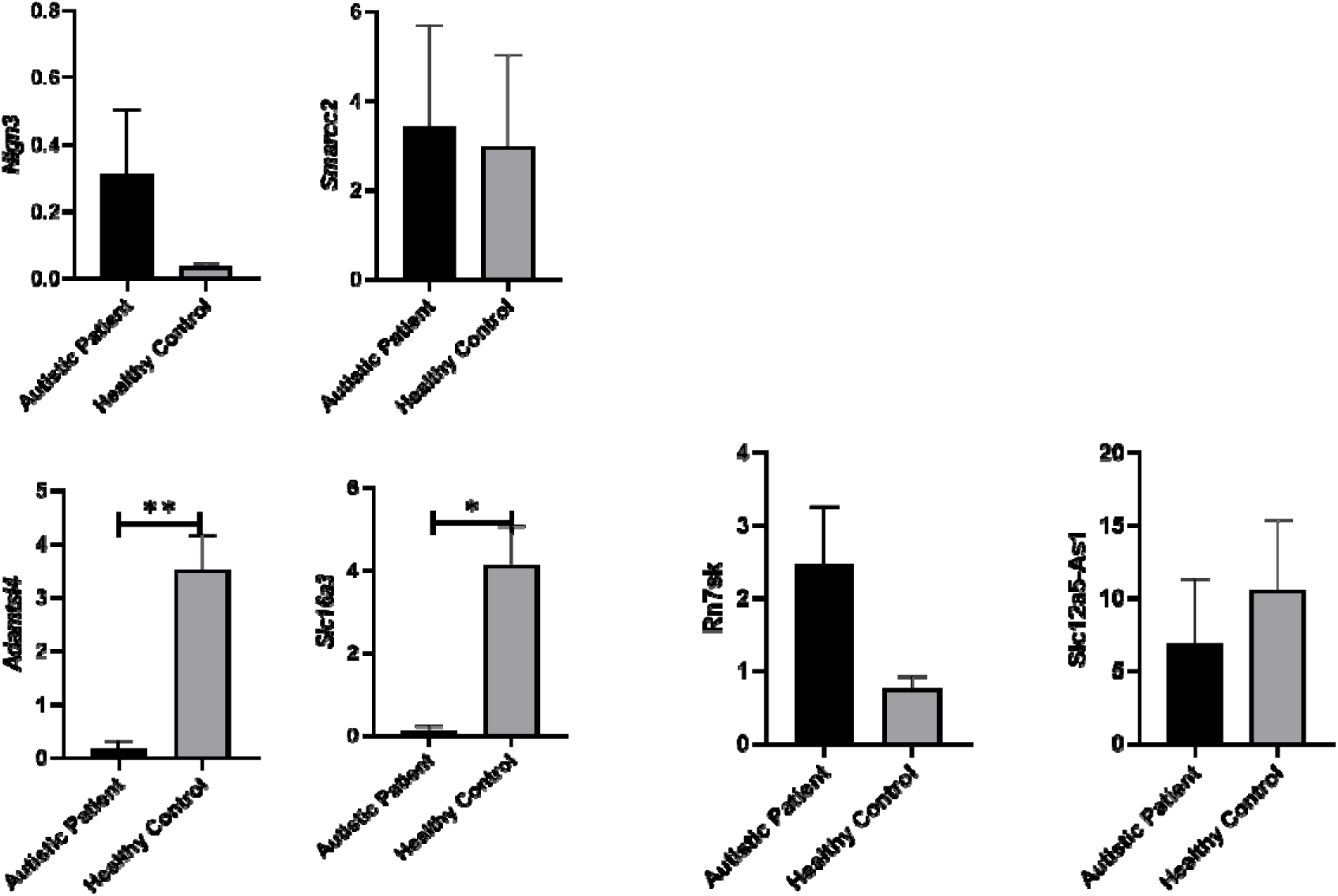
Relative expression levels of six candidate transcripts, Nlgn3, Smarcc2, Adamtsl4, Slc16a3 (protein-coding), and Rn7sk, Slc2a5-As1 (lncRNAs) in total RNAs (F) vs. GAPDH (log2). The levels in each sample were normalized to that of GAPDH (log2) (internal control). Data are presented as means± SEM. Group mean comparisons are performed using Student’s t-test.

## Discussion

The misregulation or aberrant formation of DNA/RNA hybrids may lead to transcription-associated DNA damage and genomic instability, potentially contributing to neurodevelopmental disorders. However, how the genome is dysregulated in autistic patients remains unclear. In this study, we compared the transcriptomics of DNA/RNA hybrids in ASD patients with those of healthy controls. We hypothesized that variations in hybrid regions could provide insights into the molecular mechanisms underlying autism.

To investigate whether DNA/RNA hybrid regions are altered in the genomes of ASD patients, we employed a robust method previously established by Kianmehr et al., 2019 and Rassoulzadegan et al., 2020 to recover RNA attached to genomic DNA as hybrid structures. This approach was applied to blood samples from patients with autism and healthy controls. Our findings underscore the role of non-coding RNAs in the transcriptional regulation within ASD. Non-coding RNAs are known to play a crucial role in transcriptional regulation (Mattick et al., 2023), and their dysregulation has been implicated as a potential mechanism contributing to phenotypic changes observed in various complex diseases, including autism. In a previous study, we reported decreased levels of six miRNAs in the immediate family members (parents and siblings) of patients with autism (Ozkul et al., 2020), suggesting changes at the transcriptional level.

These mechanistic insights suggest a potential autism intervention strategy for parents at risk of non-Mendelian inheritance. By assessing transcription levels in hybrid regions, it may be possible to evaluate risk factors and identify pathways that could eventually restore normal gene expression.

Recent research highlights the role of heredity and *de novo* mutations in the development of autism. While over 100 susceptibility genes or *de novo* multiplex mutations have been associated with ASD, they account for only a fraction of cases, implying that they might not be considered the cause of the disease (Cirnigliaro et al., 2023). Non-Mendelian inheritance, characterized by predominant variations in non-coding RNAs, is widely recognized as a key contributor. To our knowledge, this study provides unique insights into variations of DNA/RNA hybrid-associated non-coding RNAs as heritable signals in autism, potentially serving as markers for ASD susceptibility.

Previous studies have explored the expression of non coding RNAs in the postmortem brain of ASD patients and mouse models (Parikshak et al., 2016; Velmeshev et al., 2013) (Koç et al., 2021), suggesting their role in gene regulation and autism pathogenesis. In our study, we generated the transcriptome of DNA/RNA hybrids from blood samples, identifying over 1000 previously unannotated lncRNAs through genome-guided transcriptome assembly. Out of 1,755 transcripts selected, 1048 transcripts were classified as novel lncRNAs based on protein-coding potential.

Our assembled transcriptome yielded 278,300 transcripts, with 13,305 known transcripts expressed across all samples. Differential gene expression analysis revealed 301 transcripts significantly upregulated in ASD patients compared to healthy controls (LFC> 2, adjusted P value < 0.05). Of these, *RN7SK, SLC12A5-AS1, SLC16A3*, and *SMARCC2* were further validated by qRT-PCR. These transcripts have been confirmed to be associated with autism spectrum disorder. Notably, RN7SK and SMARCC2 showed significant differences in DNA/RNA hybrid levels between ASD samples and healthy controls. Transcripts such as *RN7SK, GIGYF1, SLC16A3*, and mitochondrial genes (*MT-ND5, MT-CYB*) showed enriched expression signals in DNA/RNA hybrids in autistic patients.

While limited information exits on RN7SK, it has been linked to neuronal cell development and differentiation (Bazi et al., 2018) as well as neuro-developmental disorders (Schneeberger et al., 2019). Interestingly, RN7SK and its pseudogenes (*RN7SKP71, RN7SKP80, RN7SKP203*), were significantly enriched in autism samples despite not being differentially expressed. Pseudogenes may regulate gene expression by acting as decoys for regulatory proteins or RNAs (Pink et al., 2011). *SMARCC2* which also showed enriched signals, encodes a subunit of the SWI/SNF chromatin remodeler and is considered a high-confidence candidates involved in ASD regulation (Nussinov et al., 2023, Ben-David and Shifman, 2013).

Gene ontology and pathway analysis revealed that upregulated genes in ASD patients were primarily involved in neurodevelopmental and neurodegenerative disorders, such as Parkinson’s, Alzheimer’s, Huntington’s, and Amyotrophic Lateral Sclerosis (ALS). Biological processes were mainly linked to mitochondrial function and cellular respiration. Despite our previous identification of miRNA-lncRNA interactions (Ozkul et al., 2020), none of the lncRNAs identified in this study were associated with the six miRNAs previously detected.

Our previous research has also highlighted variations in DNA/RNA hybrids at telomeric regions and other non-coding RNAs in multiple pathologies (Mehmetbeyoglu et al., 2022; Rassoulzadegan et al., 2020). These findings align with the emerging view that lncRNAs exhibit tissue-specific expression (Statello et al., 2021) and that structureal alterations in DNA/RNA hybrids are increasingly recognized as important contributors to human pathologies (Petermann et al., 2022). This has been demonstrated in two ASD-related conditions: Prader-Willi syndrome (PWS) and Angelman syndrome (AS), where complex epigenetic mechanisms regulate their pathology. Stabilization of DNA/RNA at GC-rich repetitive intronic regions within the Snord116 locus with a topoisomerase-I inhibitor, such as topotecan, has been shown to restore gene expression and chromatin de-condensation (Powell et al., 2013).

While these findings, along with preclinical proof-of-concept studies, are promising, more family studies are needed to understand the implications of non-Mendelian inheritance in autism. Our mechanistic understanding also calls for caution in the application of treatment, particularly when considering potential variations in gene expression.

Research into lncRNAs has been controversial due to their initially unclear functions. However, global transcriptomic analyses have unexpectedly shown that most animal and plant genomes are extensively transcribed into lncRNAs, challenging the assumption that protein-coding genes alone are responsible for driving genetic complexity. Increasing evidence suggests that lncRNAs playing pivotal roles in chromatin modification and enhancer regulation during development, emphasizing their importance in various biological processes. This study further highlights the dynamic assemblies of lncRNAs, such as RNA-DNA interactions, which have the potential to greatly expand our understanding of cell and developmental biology as well as gene-environment interactions.

A particularly innovative aspect of this study is the focus on “hybrid marks” and the exploration of pathways linking DNA to RNA. Although the exact pathogenic role of these DNA/RNA remains uncertain, resolving them promising opportunities for therapeutic interventions. The comprehensive profiling of the hybrid regions is a prerequisite for future studies, particularly in understanding how these factors might contribute to transmission of symptoms in psychiatric disorders like autism from parent to child.

Transgenerational variations in phenotypes and diseases may arise from multigenic origins or non-Mendelian heritable variations. Distinguishing between these possibilities, requires further investigation and the development of new models under controlled conditions.

### Limitations of the study

In this study, we analyzed changes in DNA/RNA hybrid fractions in blood samples from autistic patients to explore underlying pathologies. However, this analysis alone does not provide a comprehensive understanding of non-Mendelian inheritance in humans. To better elucidate the role of DNA/RNA hybrids in non-Mendelian transmission, it would be particularly valuable to examine these changes in sperm, egg, or embryo samples. Naturally, performing all such experiments on human samples presents significant challenges. As an initial step, we focused on human DNA/RNA hybrid transcriptomic data. Nevertheless, further research utilizing preclinical models, clinical studies, and experimental approaches—especially with family data—is essential for a more thorough evaluation of the role of non-Mendelian inheritance in autism.

### Conclusions

This work suggests a potential biological role for DNA/RNA hybrid variations in influencing behavior, contributing valuable insights into the molecular mechanisms underlying autism. The findings highlight how alterations in hybrid DNA/RNA regions could contribute to the pathology of autism, advancing our understanding of the disorder from a mechanistic perspective.

## Supporting information

Supplementary Files (link will be provided upon file upload)

