## Supplementary Files (link will be provided upon file upload) for "Analysis of RNA/DNA Hybrid Profiles in Blood From Autistic Patients Reveals Differences in mRNA and non-coding RNA Regions"

Table 1- Sensitivity and precision percent across multiple levels of transcripts

|  | Sensitivity | Precision |
| --- | --- | --- |
| Base level | 100% | 94.5% |
| Exon level | 93.2% | 95.6% |
| Intron level | 100% | 98.5% |
| Intron chain level | 100% | 95.6% |
| Transcript level | 99.5% | 94.2% |
| Locus level | 99.6% | 92.1% |

Table 2- List of ASD-related genes among the DEGs. The cause of the genetic association of each gene with ASD and the number of reports for each gene in relation to ASD is also present.

| **Gene symbol** | **Genetic category** | **Number of reports** |
| --- | --- | --- |
| *EPHA1* | Rare Single Gene Mutation | 6 |
| *GIGYF1* | Rare Single Gene Mutation | 14 |
| *IL1R2* | Rare Single Gene Mutation | 6 |
| *KCNJ15* | Rare Single Gene Mutation | 3 |
| *KMT2E* | Rare Single Gene Mutation, Syndromic, Genetic Association | 16 |
| *MYH10* | Rare Single Gene Mutation | 5 |
| *NLGN3* | Rare Single Gene Mutation, Genetic Association, Functional | 39 |
| *SBF1* | Rare Single Gene Mutation | 9 |
| *SGSM3* | Rare Single Gene Mutation | 6 |
| *SMARCC2* | Rare Single Gene Mutation, Syndromic, Functional | 15 |

Table 3- List of primers sequencing nucleotides forward and reverse for Real-Time PCR

| Oligo Name | 5` - Oligo Seq - 3` |
| --- | --- |
| Hsa_SLC12A5-AS1-F | CCTGAATCTGGCCACTTCGC |
| Hsa_SLC12A5-AS1-R | CTCCTTCAGTACAGGACGGC |
| Hsa_RN7SK-F | CATCCCCGATAGAGGAGGACC |
| Hsa_RN7SK-R | ATGCAGCGCCTCATTTGGATG |
| Hsa_SLC16A3_F | CCACAAGTTCTCCAGTGCCATTG |
| Hsa_SLC16A3_R | CGCCAGGATGAACACGTACATG |
| Hsa_NLGN3-F | CGGGTTGGAGTGCTAGGTTT |
| Hsa_NLGN3-R | ATATTCTCGCTCACCCAGCG |
| Hsa_SMARCC2-F | CTGTGGCTCGGCAAGAACTA |
| Hsa_SMARCC2-R | GAAATCGTAACGCCGCCATC |
| Hsa_ADAMTSL4-F | AACTACCTGGCACTTCGTGG |
| Hsa_ADAMTSL4-R | ATATCGAAAGACGGTCCCGC |

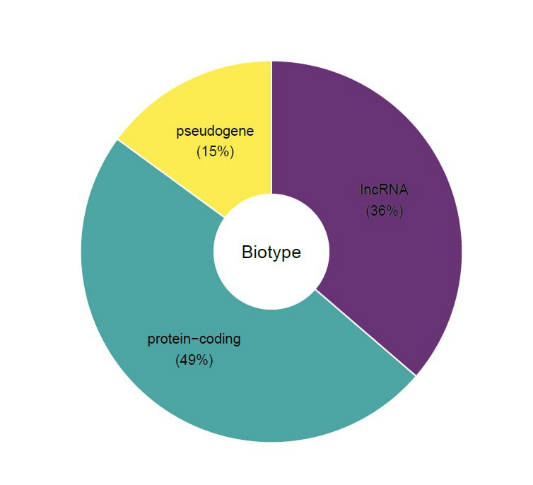

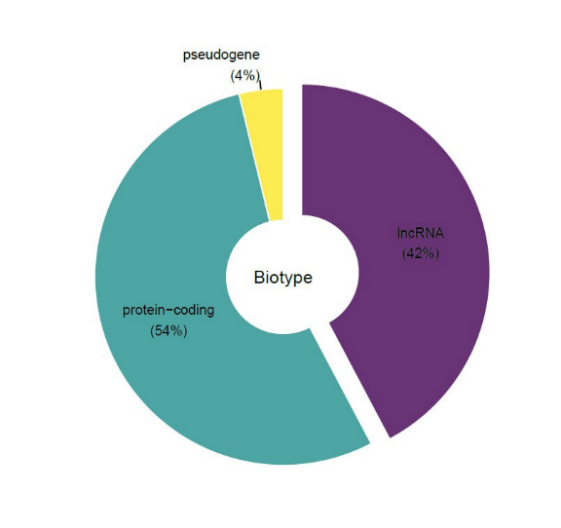

Figure 1. Types of known transcripts expressed in all blood samples. RNA types of all transcripts (left) and RNA types of differentially expressed transcripts (right).

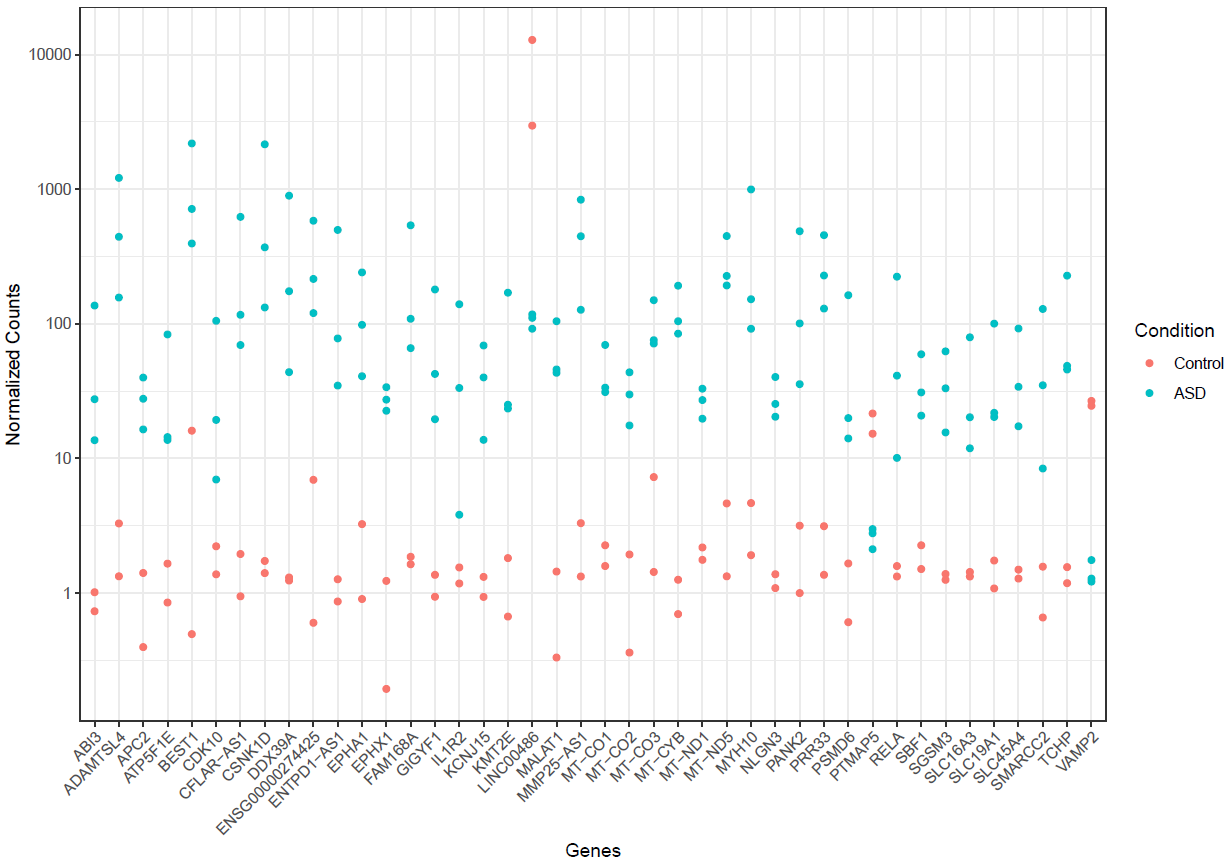

Figure 2a. The number of most significant differentially expressed transcripts in autistic patients and healthy controls. The graph plotted based on normalized counts per genes. Blue and red colors display autistic patients and healthy controls, respectively.

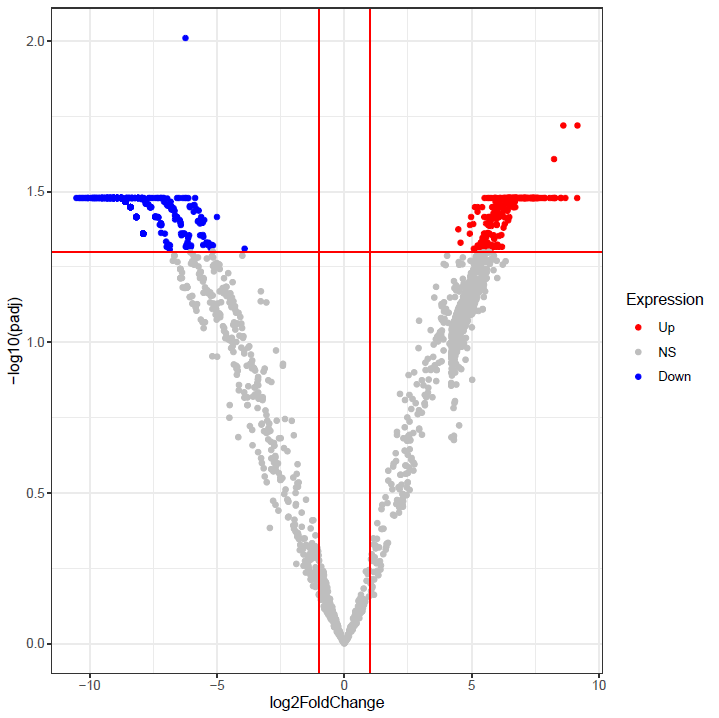

Figure 2b- Volcano plot showing differentially expressed transcripts. Plots constructed using log2 fold change and adjusted *p-*values to visualize the relationship between fold change and statistical significance. The x-axis corresponds to the fold change (log2) in transcript and the vertical axis represents the –log adjusted *p*-value. Significant differentially detected transcripts (*P*adj < 0.05) are indicated in red (higher level of transcripts in autistic patients versus healthy control log2-fold change >2) and blue (lower level of transcripts; log2-fold change < −2).

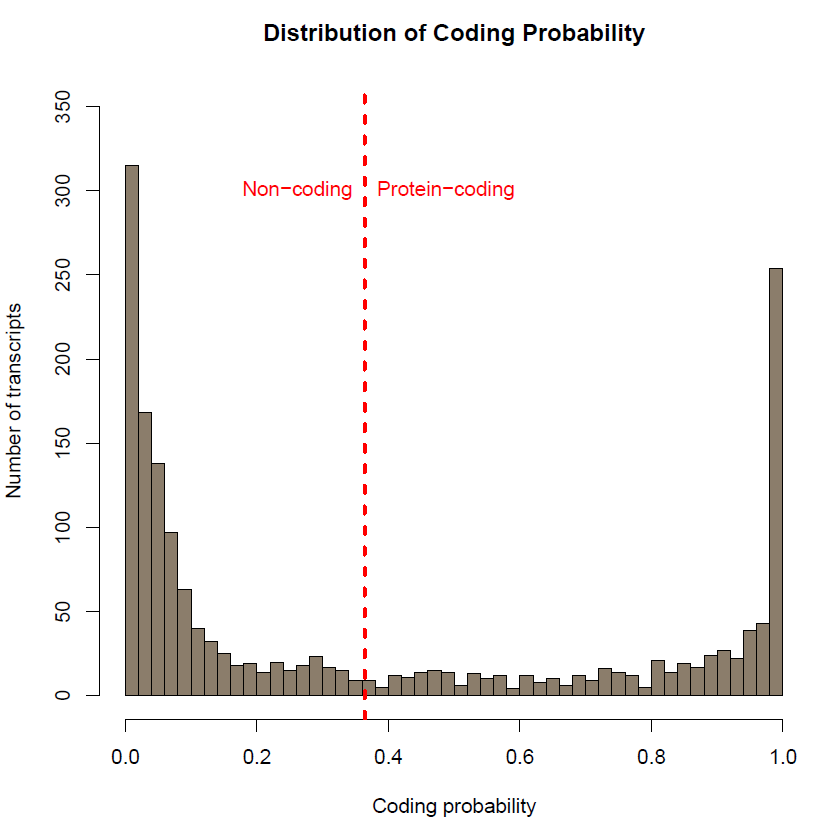

Figure 3a. Distribution of coding probability. Graph shows number of transcripts per coding probability. The dashed red line shows the threshold (0.364). Coding probability below this amount considered non-coding transcripts.

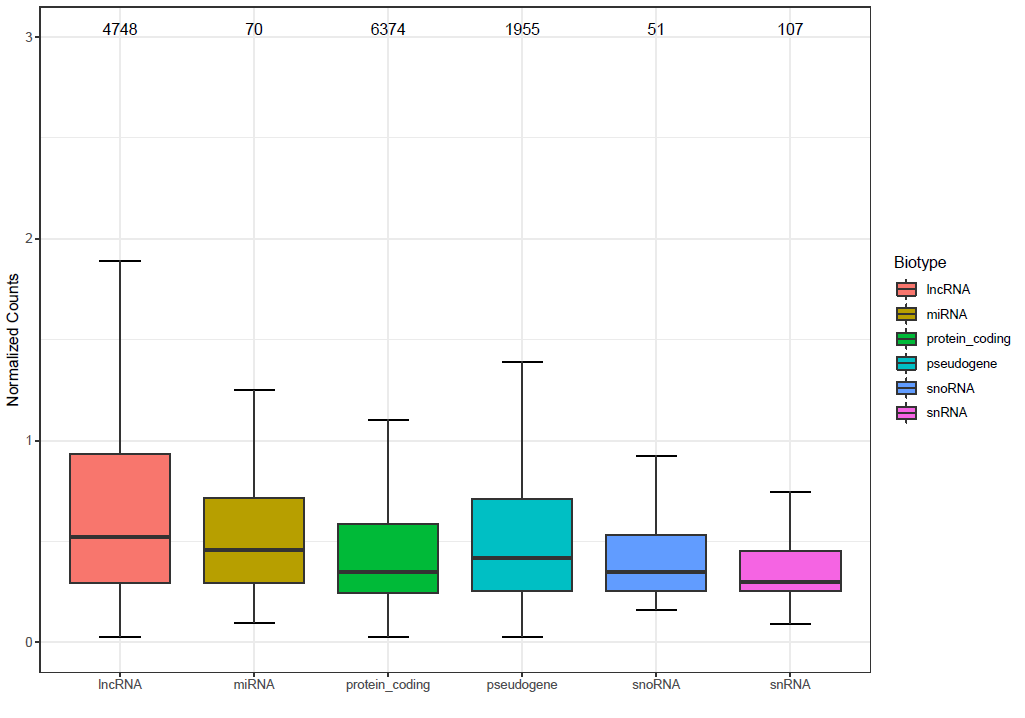

Figure3b- Box plot showing the expression level of known transcripts through all samples. Expression values are log2- normalized. The top of each bars shows the number of each transcript per genomic regions

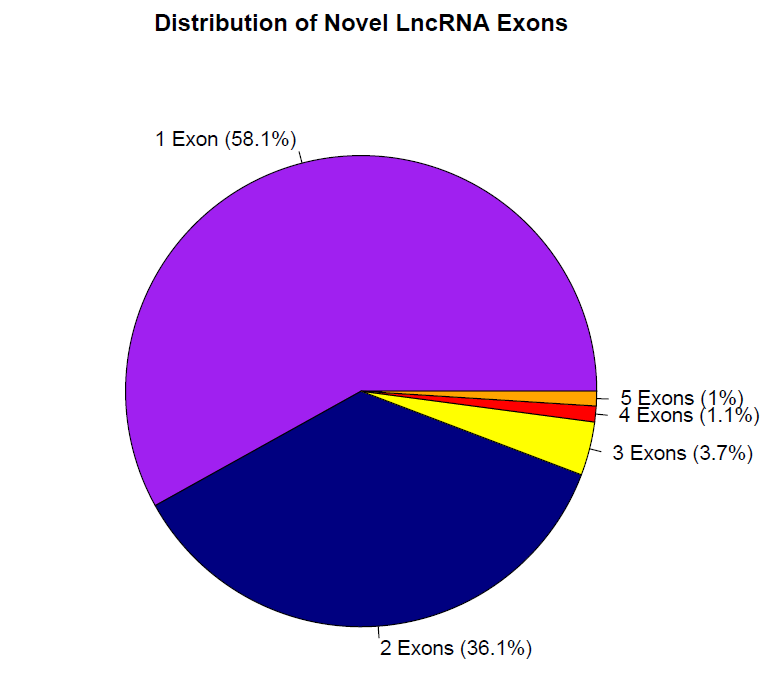

Figure 4. Exonic structure of novel lncRNAs. 94.2% of the lncRNA transcripts had either 1 or 2 exons.

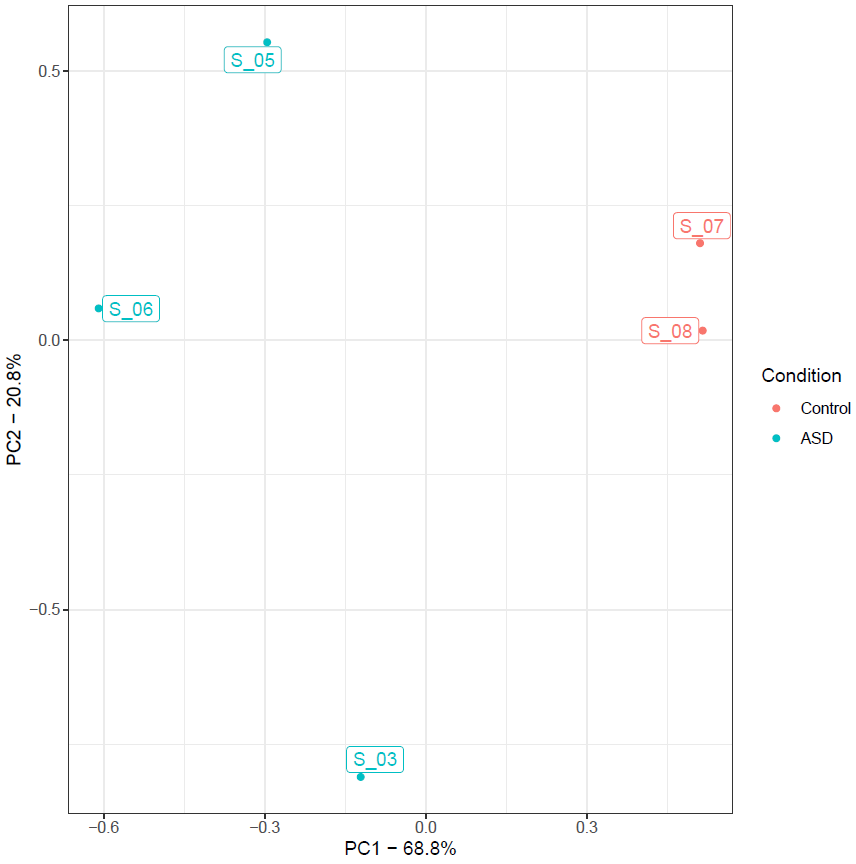

Figure 5. PCA

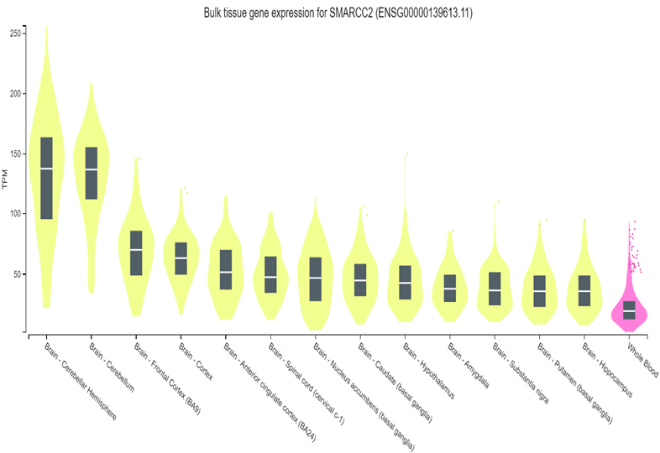

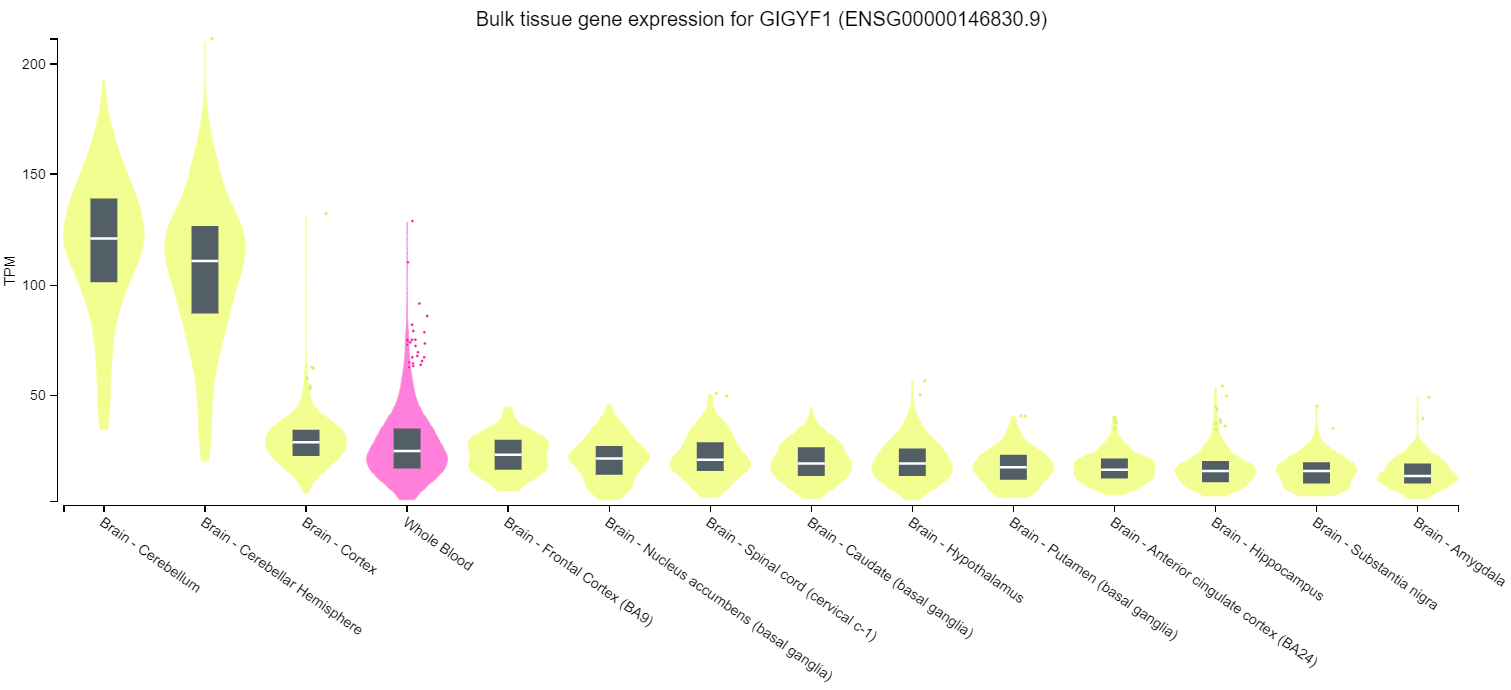

**Figure 6.**

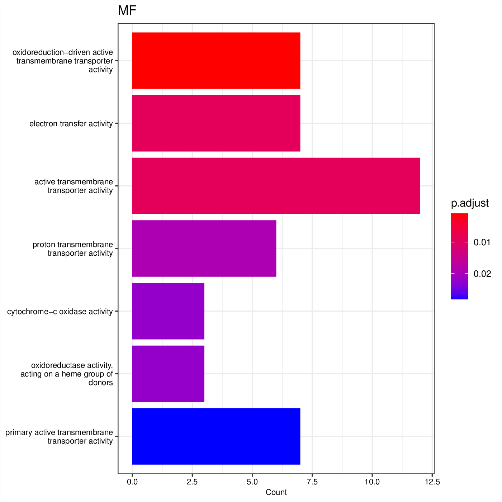

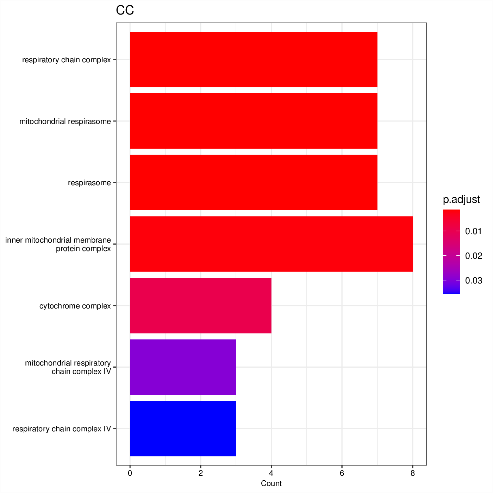

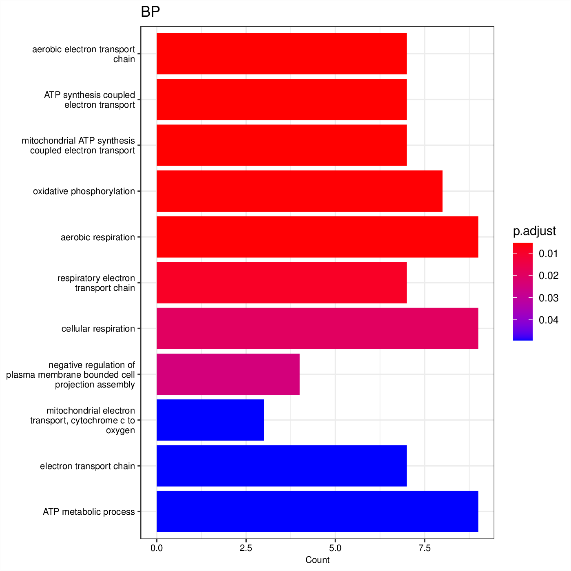

Figure 7. GO analysis of 301 up-regulated genes in ASD patients compared to healthy controls. X-axis shows the names of the GO terms. Y-axis shows the number of genes associated with each GO term. The *P-value* is adjusted by the BH method. BP: biological process, MF: molecular function, CC: cellular component.

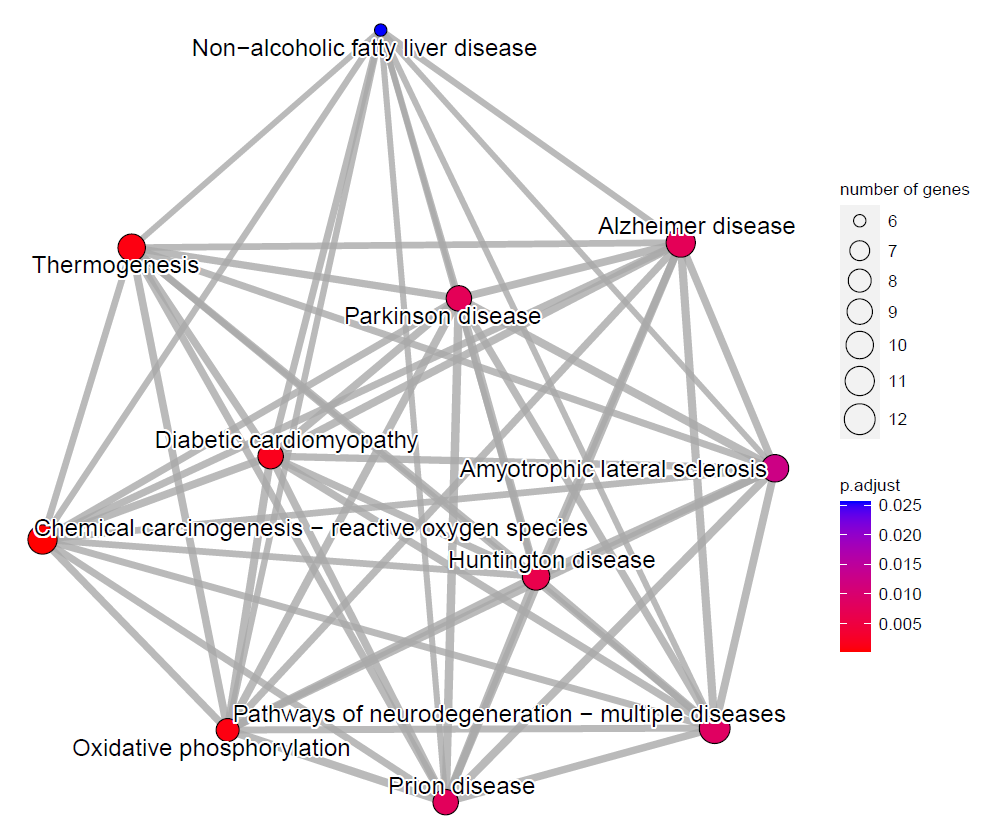

Figure 8. KEGG pathway analysis. The size of each circle reveals the number of genes involved in each GO terms. And, the scale bar displays the adjusted *p* value of each GO terms.

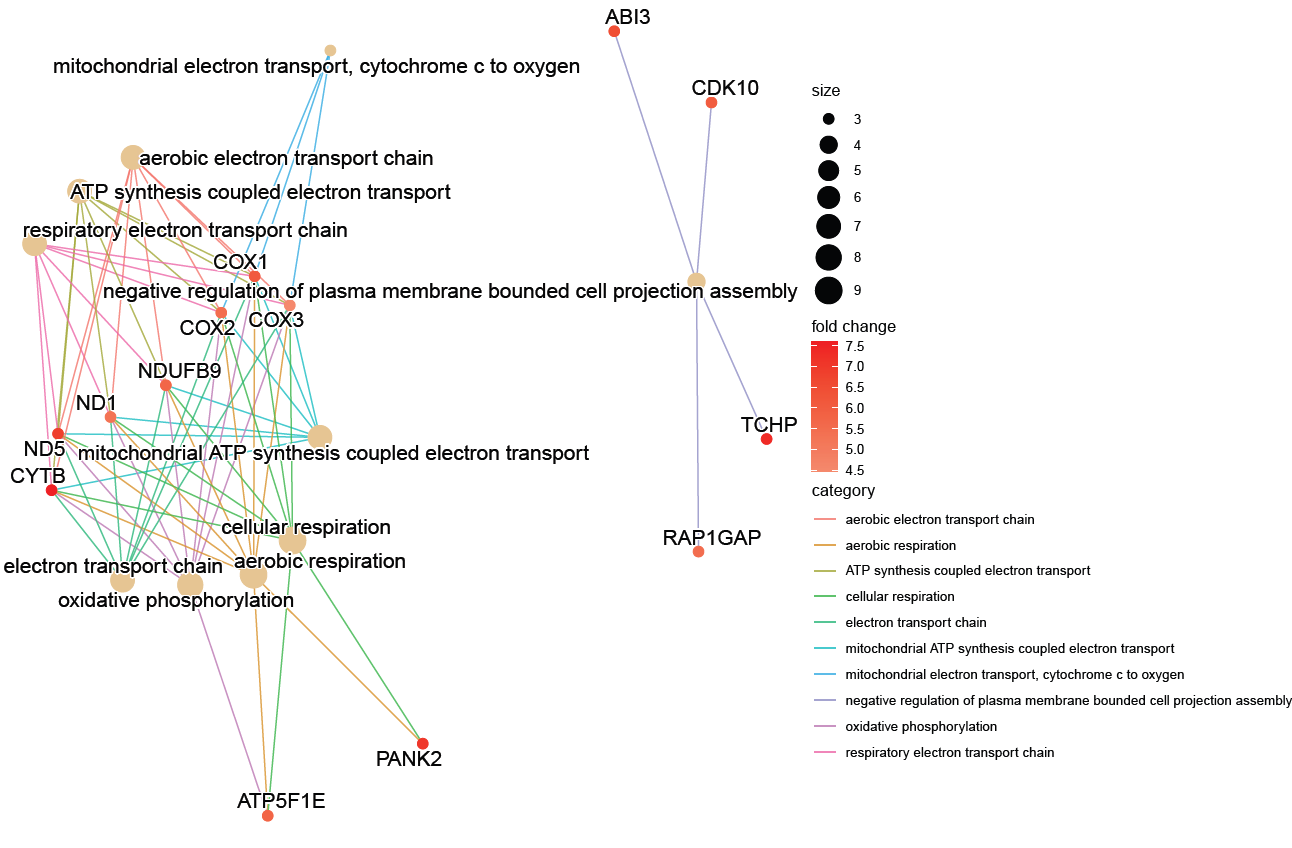

Figure 9. Genes associated with enriched GO terms in biological processes. Various genes of different mitochondrial electron transport chain complexes are present. The size of each circle representing GO terms indicates the number of genes associated with that term. Fold change (log2) is based on the normalized counts of each gene.
